# A functional genomic approach to identify reference genes for human pancreatic beta cell real-time quantitative RT-PCR analysis

**DOI:** 10.1101/2021.04.14.439798

**Authors:** Maria Inês Alvelos, Florian Szymczak, Ângela Castela, Sandra Marín-Cañas, Bianca Marmontel de Souza, Ioannis Gkantounas, Maikel Colli, Federica Fantuzzi, Cristina Cosentino, Mariana Igoillo-Esteve, Lorella Marselli, Piero Marchetti, Miriam Cnop, Décio L. Eizirik

**Affiliations:** ULB Center for Diabetes Research, Medical Faculty, Université Libre de Bruxelles, Brussels (ULB) B-1070, Belgium; Department of Clinical and Experimental Medicine, Islet Cell Laboratory, University of Pisa, Pisa, Italy; Division of Endocrinology, Erasmus Hospital, Université Libre de Bruxelles, Brussels, Belgium; Welbio, Medical Faculty, Université Libre de Bruxelles, Brussels (ULB) B-1070, Belgium; Indiana Biosciences Research Institute, Indianapolis, IN, USA

**Keywords:** Reference genes, beta cells, diabetes, RNA-sequencing, quantitative real-time PCR

## Abstract

Exposure of human pancreatic beta cells to pro-inflammatory cytokines or metabolic stressors is used to model events related to type 1 and type 2 diabetes, respectively. Quantitative real-time PCR is commonly used to quantify changes in gene expression. The selection of the most adequate reference gene(s) for gene expression normalization is an important pre-requisite to obtain accurate and reliable results. There are no universally applicable reference genes, and the human beta cell expression of commonly used reference genes can be altered by different stressors. Here we aimed to identify the most stably expressed genes in human beta cells to normalize quantitative real-time PCR gene expression.

We used comprehensive RNA-sequencing data from the human pancreatic beta cell line EndoC-βH1, human islets exposed to cytokines or the free fatty acid palmitate in order to identify the most stably expressed genes. Genes were filtered based on their level of significance (adjusted P-value >0.05), fold-change (|fold-change| <1.5) and a coefficient of variation <10%. Candidate reference genes were validated by quantitative real-time PCR in independent samples.

We identified a total of 264 genes stably expressed in EndoC-βH1 cells and human islets following cytokine- or palmitate-induced stress, displaying a low coefficient of variation. Validation by quantitative real-time PCR of the top five genes *ARF1*, *CWC15*, *RAB7A*, *SIAH1* and *VAPA* corroborated their expression stability under most of the tested conditions. Further validation in independent samples indicated that the geometric mean of *ACTB* and *VAPA* expression can be used as a reliable normalizing factor in human beta cells.

## Introduction

The study of molecular mechanisms involved in pancreatic beta cell dysfunction and death in type 1 (T1D) and type 2 diabetes (T2D) often involves *in vitro* exposure of human beta cells to stressors that may be present *in vivo* in T1D and T2D (1–3). Exposure of beta cells to these stressors, including pro-inflammatory cytokines (as a model of T1D) or palmitate (as a model of metabolic stress in T2D), substantially alters their gene expression (2, 4).

Quantitative real-time polymerase chain reaction (qPCR) is a commonly used technique to measure mRNA transcript levels owing to its sensitivity, specificity and fast execution (5). The accurate quantification of the observed changes relies on the effective normalization to one or more reference gene(s), whose expression should not be altered by the experimental condition(s) under evaluation. A unique and universal reference gene has not yet been identified, and therefore gene expression normalization usually depends on genes classified as “housekeeping genes” which, due to their cellular indispensability, are assumed to have stable expression under different experimental conditions. Commonly used reference genes, such as *beta actin (ACTB)*, *beta-2-microglobulin (B2M)*, the *18 S ribosome small subunit (18S rRNA)* and *glyceraldehyde-3-phosphate dehydrogenase (GAPDH)*, are widely used as normalizers due to their robust expression (6, 7). However, their expression varies widely among conditions and cell types (8–11), including pancreatic islets (12, 13). An inadequate selection of reference genes – that are up- or down-regulated in parallel with the gene under study - could lead to the misinterpretation of qPCR results and obscure genuine changes.

There are few studies on the identification of reference genes in human beta cells, and they are mostly limited to show that the housekeeping gene does not change under the experimental conditions used (14). Rat islets of Langerhans have different expression levels of reference genes during the first 24 hours following isolation (13), and a study in rat insulin-secreting INS-1E cells confirmed that qPCR results are affected by the normalization method and selected reference genes (15).

Selection of the most suitable reference genes should be implemented through the validation of their expression stability in the cell or tissue type under the study. Most studies use bioinformatic tools such as BestKeeper (16), geNorm (17), NormFinder (18) or Global Pattern Recognition (19), which perform a mathematical evaluation of gene expression and rank candidates according to their stability. These tools define the least variable genes from a list of pre-selected candidate genes, but they are not suitable for *de novo* identification of genes with calibrating potential.

Against this background, a genome-wide analysis to identify stable and well-expressed genes in human islets and beta cells represents an essential tool for accurate normalization. To achieve this goal, we used high-depth RNA-sequencing data from the human beta cell line EndoC-ßH1 and human islets exposed to pro-inflammatory cytokines or palmitate. Genes were validated as putative reference genes by qPCR in EndoC-βH1 cells, human islets and induced pluripotent stem cell (iPSC)-derived islets.

## Results

### *In silico* selection of new reference genes

The analysis of high-depth RNA-seq (around 200 million reads) data from insulin-producing EndoC-βH1 cells exposed to the proinflammatory cytokines interleukin-1β (IL1β) plus interferon-γ (IL1β + IFNγ) for 48h, or interferon-α (IFNα) for 2h, 8h, and 24h and human islets exposed to IL1 β + IFNγ for 48h or IFNα for 2h, 8h, and 18h, or the metabolic stressor palmitate for 48h (**Figure 1A**) (see references in Methods) indicated that 17,874 ± 3,751 genes are not significantly modified, according to the criteria of an adjusted P-value >0.05 and |fold-change| <1.5. The gene expression level quantified as transcripts per million (TPM) was used to evaluate intra- and inter-experimental condition variability, and 264 genes display a coefficient of variation (CV) <10%, of which 175 genes have a mean TPM between 10 and 100, and 89 a mean TPM between 100 and 1000 (**Figure 1B-D**) (**Supplementary Table 2**). Genes with mean TPM >1000 (**Figure 1B, C, D**) have the highest mean CV. For this reason, we selected the candidate reference genes among the low (10<mean TPM<100) and intermediate TPM (100<mean TPM<1000) intervals. Gene Ontology (GO) enrichment analysis on this set of 264 genes revealed an enrichment in pathways associated with central cellular functions such as “intracellular transport”, “mRNA processing”, “RNA splicing”, and “protein catabolic processes” (adjusted P-value <0.05) (**Figure 1E**). As a proof of concept, we selected five genes: *ADP-ribosylation factor 1 (ARF1)*, *spliceosome associated protein homolog (CWC15)*, *member RAS oncogene family (RAB7A)*, *E3 ubiquitin protein ligase 1 (SIAH1)* and *vesicle-associated membrane protein-associated protein A (VAPA)* from the 264 candidates for subsequent validation. Except for *RAB7A* (20), none of the genes have been previously used as reference genes in human beta cells or other tissues. The mean CV for the selected 5 genes ranged from 7% to 8.4%, and they were more stably expressed in human beta cells as compared to *ACTB* and *GAPDH*, which have CVs of 17.9% and 23.4%, respectively (**Figure 1C, D**). Interestingly, *ARF1*, *CWC15*, *RAB7A*, *SIAH1*, *VAPA* genes have the same mean TPM distribution and CV in beta cells from T1D and T2D donors and non-diabetic controls, reinforcing our observation that these genes are stably expressed, and may be suitable human beta cell reference genes (**Supplementary Figure 1A, B**).

**Figure 1.**
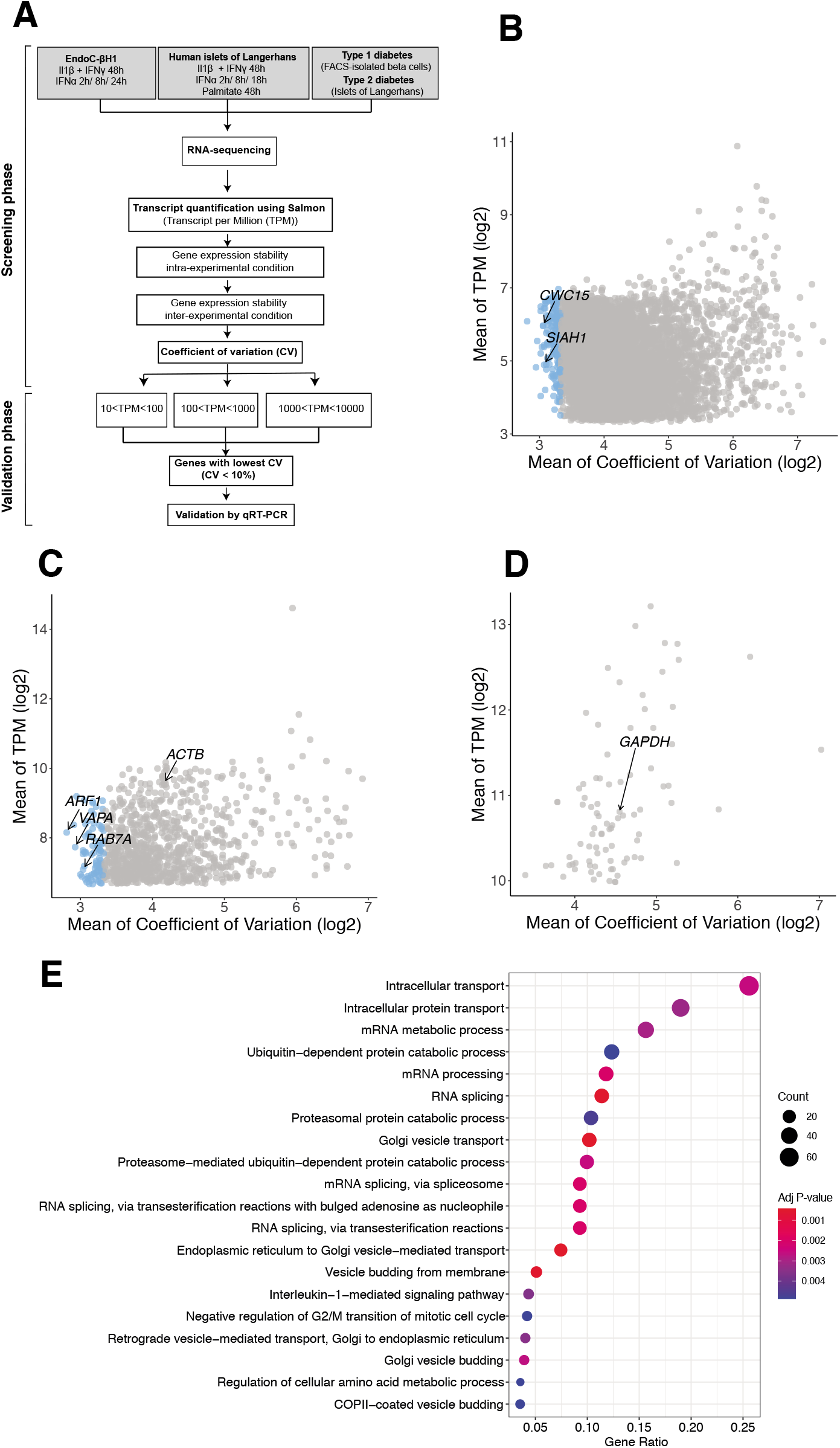
Identification of new reference genes for human islets/beta cells. **(A)** The approach for the identification of new reference genes in human pancreatic beta cells started with the analysis of RNA-sequencing data from EndoC-βH1 cells exposed to IL1β + IFNγ (48h, n=5) or IFNα (2h, 8h, and 24h, n=5 per time point); human islets exposed to IL1β + IFNγ (48h, n=5) or IFNα (2h, 8h, and 18h, n=6 per time point); human islets exposed to palmitate (48h, n= 5); fluorescence-activated cell sorting (FACS)-purified beta cells from T1D patients (n=4), and islets from T2D patients (n=28). The quantification of expression of gene transcripts was conducted using Salmon v1.3.0 (Genome reference: GENCODE GRCh38.p13) (58). Transcript expression levels, measured in transcripts per million (TPM), were used to calculate the intra- and inter-sample stability, and their respective coefficient of variation (CV). TPM values were subdivided in three categories: 10<TPM<100; 100<TPM<1000; 1000<TPM<10,000. Genes with the lowest mean CV were selected for further validation by qRT-PCR. **(B,C,D)** Scatterplots of mean CV against mean expression values (TPM) after log2 transformation based on RNA-seq data, for genes with TPM values between 10 and 100 **(B)**, between 100 and 1000 **(C)**, and between 1000 and 10,000 **(D)**. Each circle represents a gene with an adjusted P-value >0.05, |fold-change| <1.5, and blue circles represent the genes with CV <10%. Selected reference genes are shown in the scatterplots, along with the well-established reference genes *ACTB* and *GAPDH*. **(E)** Significantly enriched Gene Ontology (GO) terms by gene enrichment analysis of the 264 genes with adjusted P-value >0.05, |foldchange <1.5| and CV <10%. The gene ratio represents the ratio between the genes in the GO term that overlaps with the query gene list (the short list of candidate genes with CV <10%). The top 20 enriched GO terms are represented. Adjusted P-value from hypergeometric distribution (Benjamini-Hochberg).

### Validation of gene expression stability by quantitative real time PCR (qPCR)

In order to assess the validity of the RNA-seq findings, we quantified by qPCR the expression of the five selected potential reference genes (*ARF1*, *CWC15*, *RAB7A*, *SIAH1*, *VAPA*) and the commonly used (*ACTB* and *GAPDH*) reference genes in independent samples of EndoC-βH1 cells and human islets exposed to different stress conditions. To allow accurate comparison between samples, all of them were diluted to the same final cDNA concentration, as described in Methods. PCR amplification efficiency was >90% for all genes.

The exposure of EndoC-βH1 cells to IL1β + IFNγ or IL1β + IFNα did not modify the expression of the analyzed reference genes (**Figure 2A, B**). However, IFNα modified expression of several of them, including *ACTB* and *GAPDH* (**Figure 2C, D**). Although *ACTB* and *GAPDH* have higher CV values compared to the novel reference genes in the RNA-seq data (Figures 1C and D), qPCR of these genes indicated stable expression in most conditions (**Figure 2, 3, 4**, **Supplementary Figure 1**) except for IFNα exposure. A combination of reference genes can provide more accurate correction for mRNA loading (21). We selected *ACTB* and *VAPA* because they are amongst the most well-expressed genes, displaying quantification cycle (Cq) values between 17-28 and 20-28, respectively, do not change following exposure to different stresses (**Figure 1**) and, at least for *ACTB*, are already in use by many islet research laboratories. We calculated the geometric mean between *ACTB* and *VAPA* Cq values and included them for comparison in the figures.

**Figure 2.**
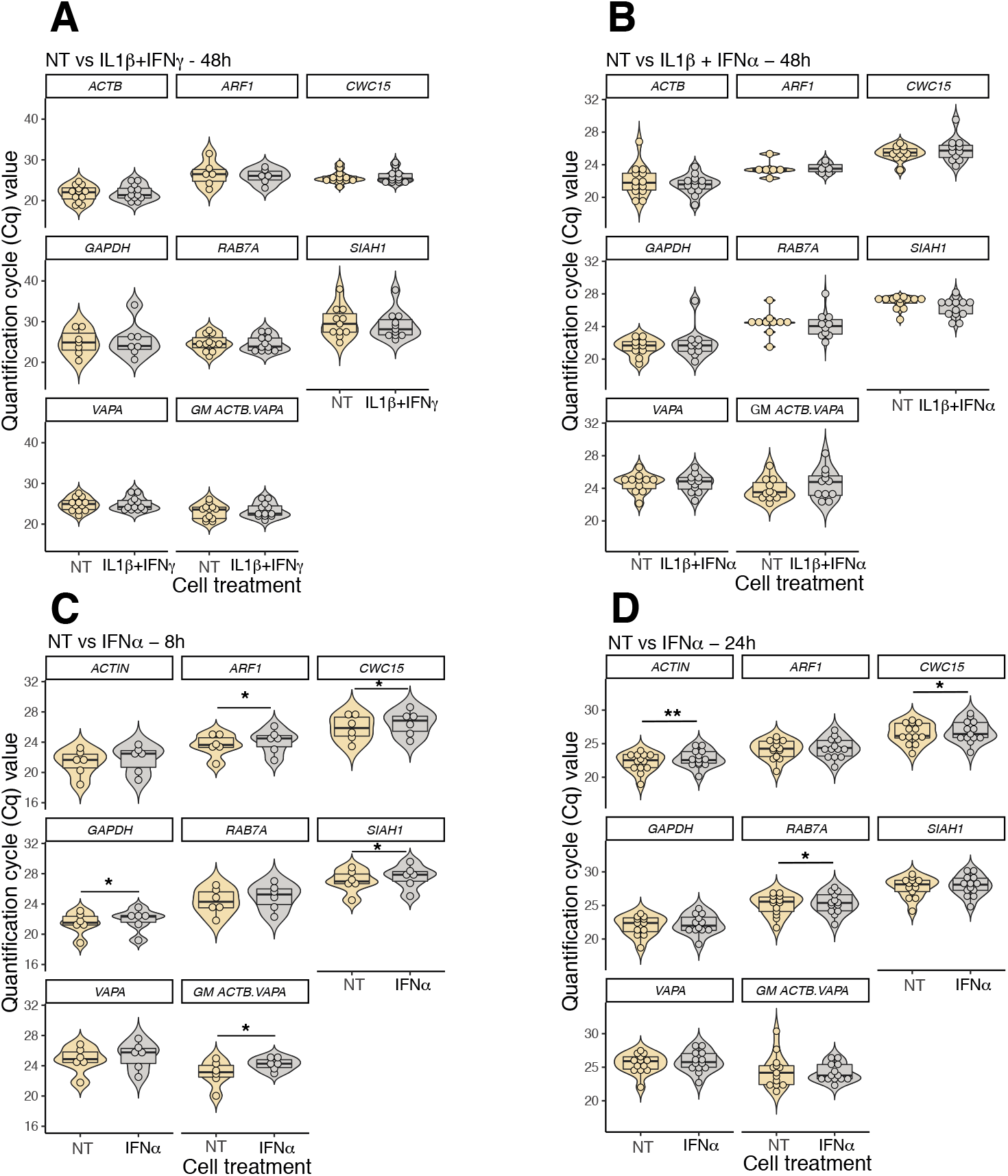
Validation of reference genes by qPCR in EndoC-βH1 cells. Distribution of the quantification cycle (Cq) value of five new (*ARF1*, *CWC15*, *RAB7A*, *SIAH1*, *VAPA*) and two commonly used reference genes (*ACTB* and *GAPDH*) based on qRT-PCR, in EndoC-βH1 cells treated with IL1β + IFNγ for 48h **(A)**, IL1β + IFNα for 48h **(B)**, IFNα for 8h **(C)** and 24h **(D)**. Geometric means of *ACTB* and *VAPA* Cq values were also calculated (GM *ACTB VAPA*). The shape of the violin plots reflects the distribution of the data, and the width of curve corresponds to the frequency of values in each region. The integrated boxplots show the 25^th^ and 75^th^ percentiles, and the horizontal line represents the median. Results from six to thirteen independent experiments. *P <0.05, **P <0.01 *versus* non-treated (NT) (paired Student’s t-test).

**Figure 3.**
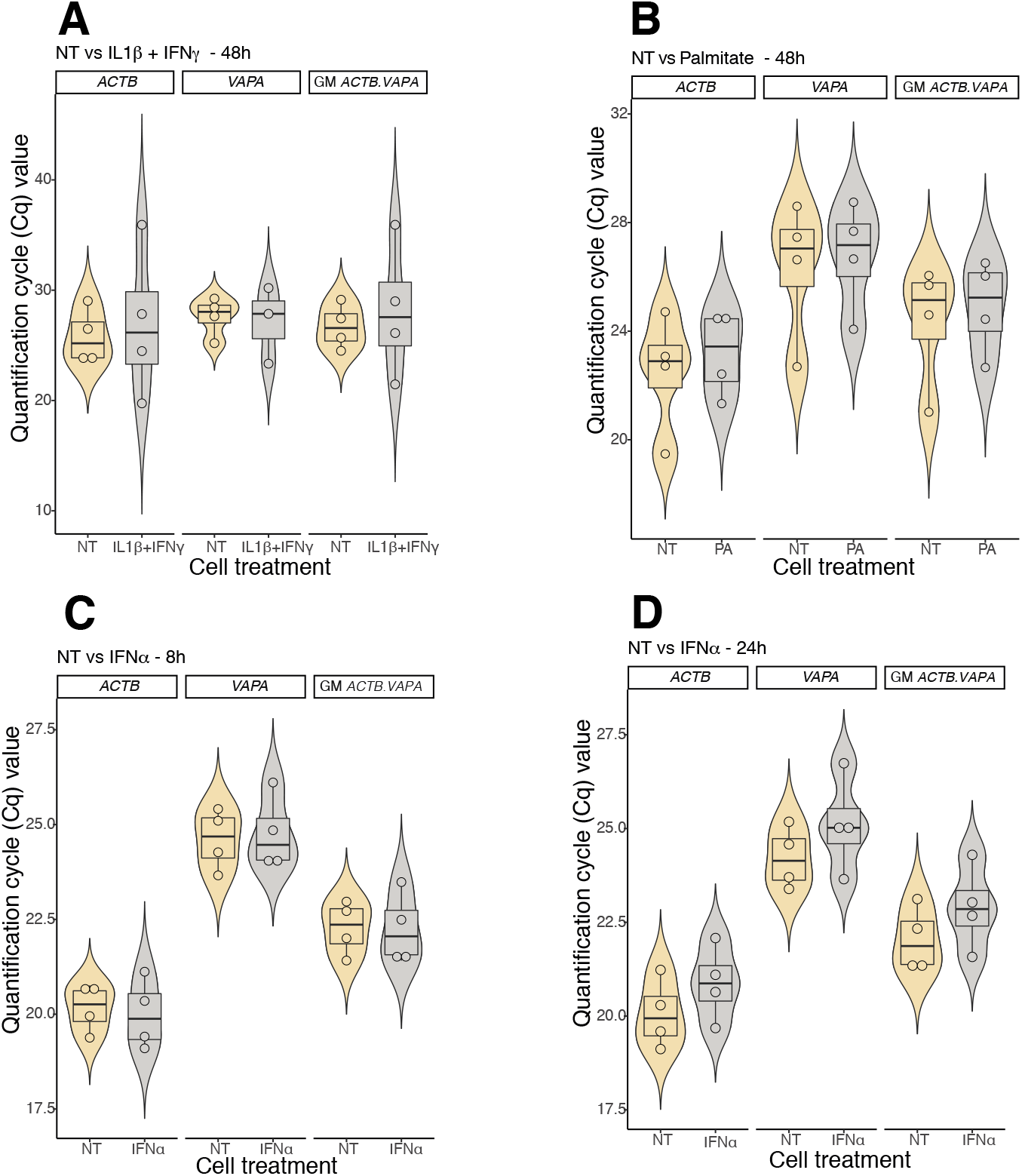
Validation of reference genes by qPCR in dispersed human islet cells. Distribution of the quantification cycle (Cq) value of five new and two commonly used reference genes (*ACTB* and *GAPDH*) based on qRT-PCR, in human islet cells treated with IL1 β + IFNγ for 48h **(A)**, palmitate for 48h **(B)**, IFNα for 8h **(C)** and 24h **(D)**. Geometric means of *ACTB* and *VAPA* Cq values were also calculated (GM *ACTB VAPA*). The violin plots’ shape reflects the distribution of the data, and the width of curve corresponds to the frequency of values in each region. The integrated boxplots show the 25^th^ and 75^th^ percentiles, and the horizontal line represents the median. Results from four independent experiments. There is no statistical difference between the tested conditions (paired Student’s t-test, condition *versus* non-treated (NT)).

**Figure 4.**
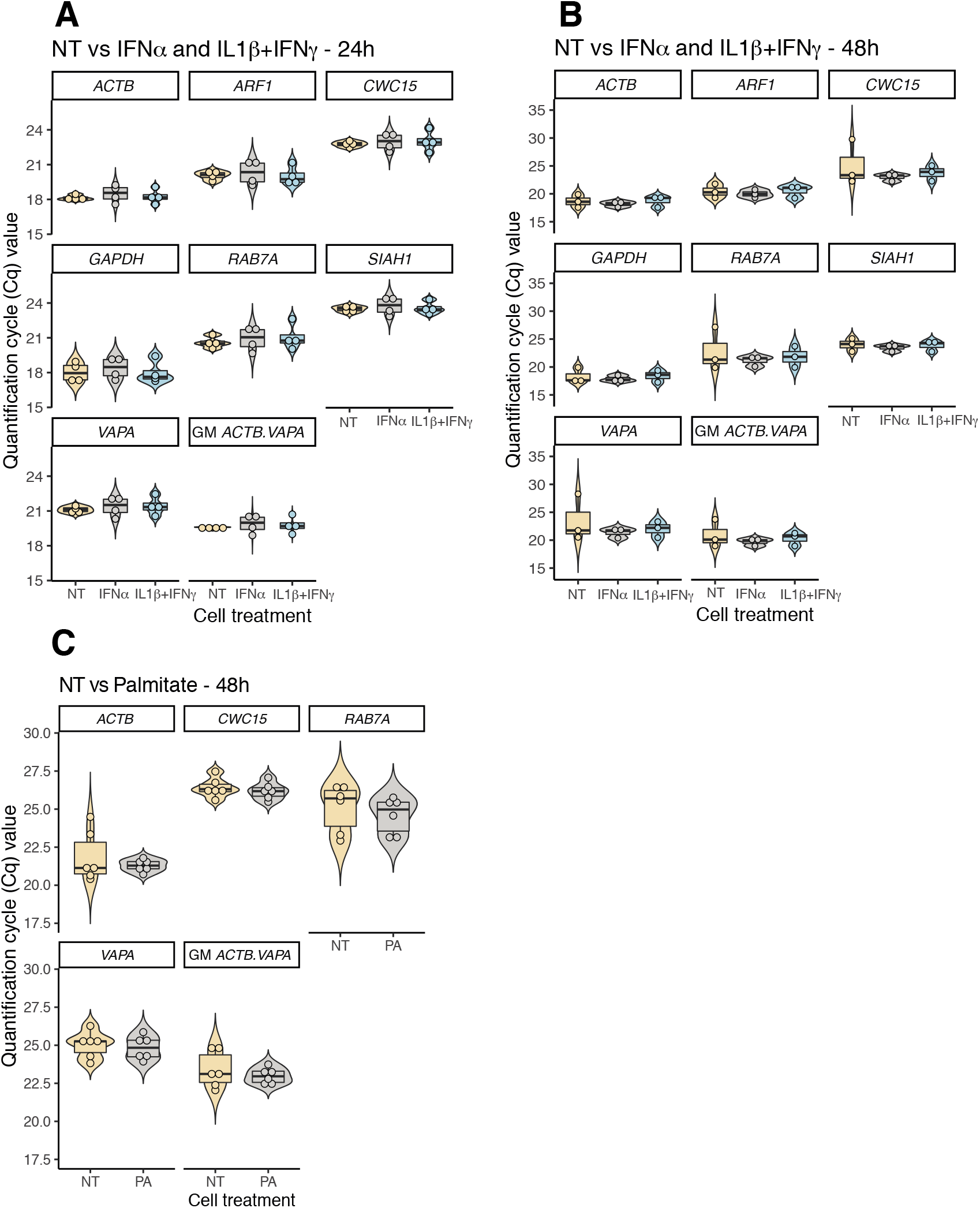
Validation of reference genes by qPCR in human iPSC-derived beta cells. Distribution of the quantification cycle (Cq) value of five new and two commonly used reference genes (*ACTB* and *GAPDH*) based on qPCR of human iPSC-derived beta cells treated with IL1β + IFNγ and IFNα for 24h **(A)**, IL1β + IFNγ and IFNα for 48h **(B)**, and the metabolic stressor palmitate for 48h **(C)**. Geometric means of *ACTB* and *VAPA* Cq values were also calculated (GM *ACTB VAPA*). The violin plots’ shape reflects the distribution of the data, and the width of curve corresponds to the frequency of values in each region. The integrated boxplots show the 25^th^ and 75^th^ percentiles, and the horizontal line represents the median. Results from four to six independent experiments. There is no statistical difference between the tested conditions (paired Student’s t-test, condition *versus* non-treated (NT)).

We next tested the effect of thapsigargin (TG) and tunicamycin (TM), two commonly used chemical endoplasmic reticulum stressors. EndoC-βH1 cells exposed to TG for 24h or TM for 48h neither showed differences in Cq values of the five new reference genes nor on *ACTB* and *GAPDH* expression (**Supplementary Figure 1C, D**).

Due to the limited amount of human islet material available, we restricted our analysis to *ACTB* and *VAPA*. IL1β + IFNγ, IFNα or palmitate did not change expression of *ACTB* and *VAPA* genes in human islets (**Figure 3**).

iPSC-derived beta cells represent a new model to study pathophysiologic mechanisms in T1D and T2D (22–27). We analyzed candidate reference genes in iPSC-derived islets exposed to the pro-inflammatory cytokines IL1β + IFNγ and IFNα and palmitate (**Figure 4A, B, C**). None of these stressors altered the expression of the genes, suggesting that they are suitable for qPCR data normalization in iPSC-derived islets.

### Validation of the necessity for additional reference genes

To validate the normalization strategies for genes of particular interest in the context of islet inflammation, we compared mRNA expression of *IRF1* (a key transcription factor downstream of IFNα signaling in beta cells (28–30)) before and after IRF1 knockdown in EndoC-βH1 cells under basal conditions and following IFNα exposure. Normalization of IRF1 expression by *ACTB* alone failed to show *IRF1* silencing following transfection with *IRF1* siRNA (**Figures 5A**). Normalization by the geometric mean of *ACTB* and *VAPA* unveiled a significant *IRF1* inhibition (**Figure 5B**), which is probably due to the reduced CV in all four conditions analyzed (**Figure 5C**). Similarly, mRNA induction of *HLA-ABC* (a key component of islet antigen presentation in the context of T1D (31)) by IFNα (**Figure 5D, E, F**) was detected following normalization by the geometric mean of *VAPA* and *ACTB*, with several fold decreased CV (**Figure 5F**).

**Figure 5.**
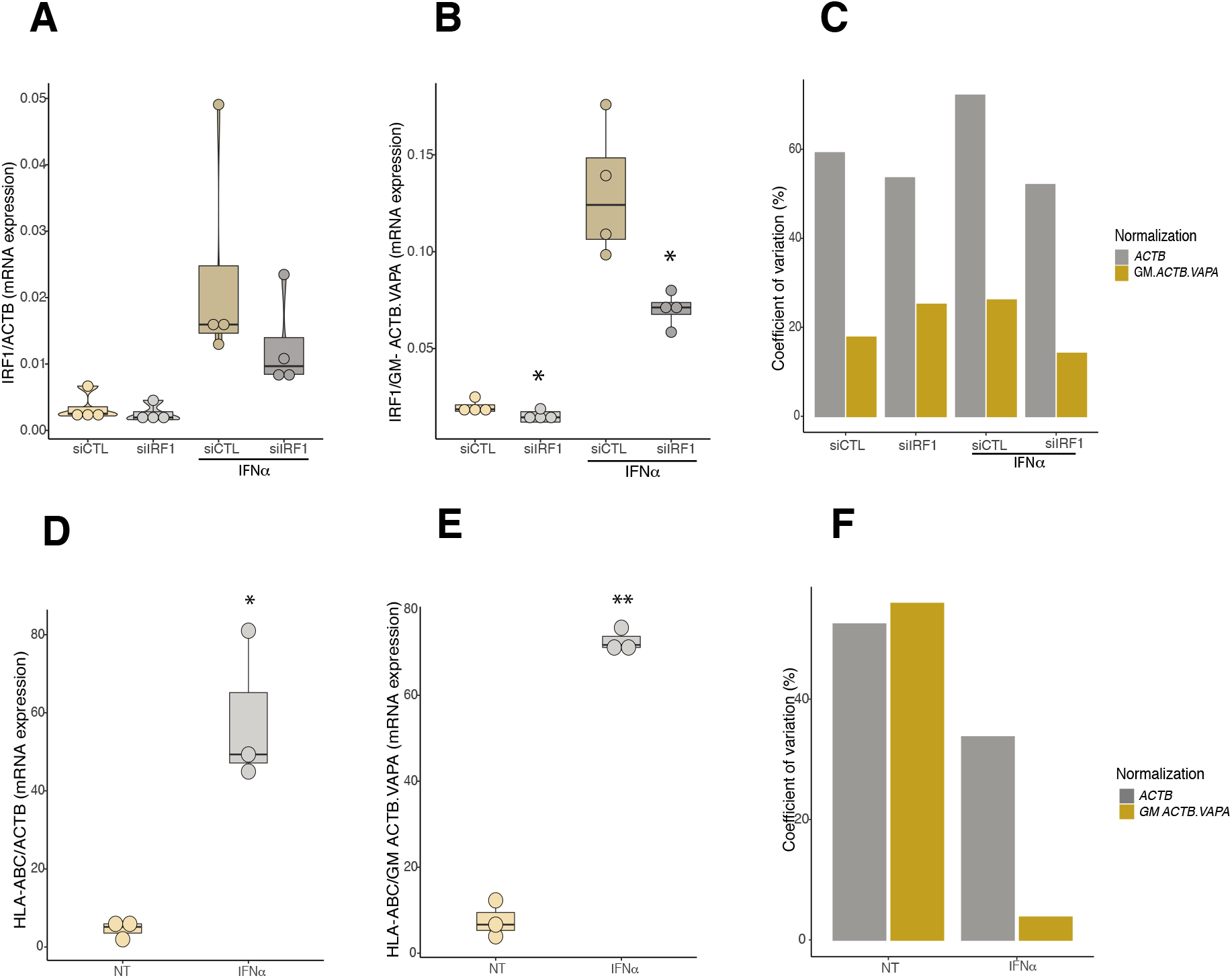
Quantification of *IRF1* and *HLA-ABC* mRNA expression in EndoC-βH1 cells. EndoC-βH1 cells were transfected with control siRNA (siCTL) or siRNA against IRF1 (siIRF1), for 48h, and then treated with IFNα for 24h. IRF1 mRNA expression was normalized to *ACTB* **(A)**, and to the geometric mean of *ACTB* and *VAPA* **(B)**. Normalization by the geometric mean of *ACTB* and *VAPA* decreases IRF1 expression variability (lower CV) **(C)**. EndoC-βH1 cells were treated with IFNα for 24h and HLA-ABC expression was normalized to ACTB **(D)**, and to the geometric mean of ACTB and VAPA **(E)**. Normalization by the geometric mean of *ACTB* and *VAPA* decreases HLA-ABC expression variability (lower CV) **(F)**. The boxplots show the 25^th^ and 75^th^ percentiles, and the horizontal line represents the median. Results from four independent experiments. *P <0.05, **P <0.01 *versus* siCTL (paired Student’s t-test).

## Discussion

We presently combined different strategies to identify new reference genes for qPCR analysis in human pancreatic beta cells. We used RNA-seq data from beta cells or human islets exposed to different stresses and identified 264 top stably expressed genes. These genes are mostly related with cellular housekeeping functions, as indicated by the fact that they are enriched in GO biological pathways related to intracellular (protein) transport, mRNA metabolic processes etc (**Figure 1E**). Following qPCR validation in independent samples, we found that the 5 candidate reference genes *ARF1*, *CWC15*, *RAB7A*, *SIAH1* and *VAPA* fit the criteria of ‘suitable reference genes’ (21). These genes have stable transcript abundance under all biologic contexts tested and are, in general, comparable to the expression of target genes. Since the geometrical mean of two or more reference genes provides a more accurate correction (17), we selected *VAPA* and *ACTB* as a suitable compromise between a novel, highly stable mRNA, and a well-established but more variable reference mRNA.

Our data indicate that the geometric mean of *ACTB* and *VAPA* expression represents a valid normalization strategy, as these gene are stably expressed (present data) and their combined use reduces the CV of target genes such as IRF1 and HLA class I (**Figure 5**). qPCR guidelines suggest to use at least 3 reference genes (21), but taking into account the scarcity of human islets, and the present data showing that it is possible to obtain reliable data using only two reference genes, we suggest that the geometrical mean of *ACTB* and *VAPA* provides a fair compromise.

*ACTB* encodes one of six different actin isoforms, and has been widely used as a ubiquitously expressed reference gene (32). Actin-B protein is highly conserved, and maintains cell structure, integrity, and motility (33, 34). Its rate of transcription is affected by mitogenic stimuli such as epidermal growth factor, transforming growth factor-β (TGF-β) and platelet derived growth factor (35–37). Since human beta cells proliferate very little (38), this should not present a major issue when using *ACTB* as reference gene.

*VAPA* encodes a protein involved in the formation of endoplasmic reticulum contacts with other membranes, such as the Golgi complex and endosomes (39, 40). It participates in fundamental physiological processes such as vesicle trafficking, membrane fusion, protein complex assembly and cell motility (41, 42). The role of *VAPA* is consistent with housekeeping functions and its expression does not change under the presently tested experimental conditions. Of particular relevance, VAPA expression is not changed in beta cells from T1D patients or whole islets from T2D patients.

In conclusion, the present study identified a panel of genes that can be used as reference for qPCR studies in human beta cells (Supplementary Table 1). The geometrical mean of two of these, namely *ACTB* and *VAPA*, may provide a robust normalization tool in the study of human beta cells. It will be important to confirm in different experimental conditions that neither *ACTB* nor *VAPA* are changed. Should this be the case, the other presently identified reference genes may be tested as an alternative approach.

## Material and methods

### Culture of human cells, gene silencing and treatments

The human beta cell line EndoC-βH1 was provided by Dr R. Scharfmann (Institut Cochin, Université Paris Descartes, Paris, France) (43). EndoC-βH1 cells were cultured in Dulbecco’s Modified Eagle Medium (DMEM) containing 5.6 mmol/L glucose (Gibco, Thermo Fisher Scientific), 2% fatty acid-free bovine serum albumin (BSA) fraction V (Roche), 50 μmol/L 2-mercaptoethanol (Sigma-Aldrich), 10 mmol/L nicotinamide (Calbiochem), 5.5 μg/mL transferrin (Sigma-Aldrich), 6.7 ng/mL selenite (Sigma-Aldrich), 100 U/mL penicillin + 100 μg/mL streptomycin (Lonza) in matrigel–fibronectin–coated plates (44).

Human islets were isolated from non-diabetic organ donors by collagenase digestion and density gradient purification and characterized as previously reported (45), with the approval of the the local Ethical Committee in Pisa, Italy. After isolation, the islets were cultured in M199 culture medium (5.5 mmol/L glucose) and shipped to our laboratory. On arrival, the islets were cultured in Ham’s F-10 medium containing 6.1 mmol/L glucose (Gibco, Thermo-Fisher Scientific), 10% fetal bovine serum (Gibco, Thermo-Fisher Scientific), 2 mmol/L GlutaMAX (Sigma-Aldrich), 50 mmol/L 3-isobutyl-1-methylxanthine (Sigma-Aldrich), 1% fatty acid-free BSA fraction V, 50 U/mL penicillin and 50 mg/mL streptomycin (44).

The iPSC lines HEL46.11 (46) and HEL115.6 (22) were derived from human neonatal foreskin and umbilical cord fibroblasts, respectively. The iPSC line Wolf2010-9 was kindly provided by Dr. Fumihiko Urano (Washington University School of Medicine, St. Louis, MO, USA) and it is derived from patients with Wolfram syndrome (47). iPSCs were cultured in E8 medium (Life Technologies) in matrigel-coated plates (Corning BV, Life Sciences). iPSCs were differentiated into pancreatic beta cells using a 7-step protocol as previously described (22, 46, 48, 49).

EndoC-βH1 cells, dispersed human islet cells and iPSC-beta cells were exposed to human pro-inflammatory cytokines IL1β (50 U/mL; R&D Systems) and IFNγ (1,000 U/mL; PeproTech) for 24h and/or 48 h, as described (2, 23). EndoC-βH1 cells, dispersed pancreatic islets and iPSC-beta cells were exposed to human IFNα (2000 U/mL; PeproTech) alone for 2h/8h/24h, 2h/8h/18h and 24-48h, respectively (23, 50, 51). EndoC-βH1 cells were also exposed to a combination of IFNα and IL1β (50 U/mL; R&D Systems) for 48h (50, 51). These conditions are based on previously published time course and dose-response experiments (44, 50).

Dispersed human islets and iPSC-beta cells were exposed to 0.5 mmol/L palmitate (Sigma-Aldrich) for 24 h and 48h, respectively (1, 4, 52).

EndoC-βH1 cells were exposed to thapsigargin (1 μM; Millipore-Sigma) and tunicamycin (5 μg/mL; Sigma-Aldrich) for 24h and 48h, respectively (53, 54).

### RNA-sequencing

The RNA-seq experiments of EndoC-βH1 cells, dispersed human islets, and islets from T2D patients were previously published by our group (30, 45, 55). RNA-seq data of FACS-purified beta cells from T1D patients was downloaded via the Gene Expression Omnibus (GEO) repository (56). All RNA-seq data were re-analyzed using our own and up-dated pipeline. Initial quality control of reads was assessed using FastQC (version 0.11.5; FastQC: A quality control tool for high throughput sequence data [Online]. Available at: http://www.bioinformatics.babraham.ac.uk/projects/fastqc/) and gene expression was quantified (in TPM) using Salmon v1.3.0(57) with extra parameters “--seqBias –gcBias --validateMappings”. The reference genome (GENCODE gene annotation release 31(58)) was indexed using default parameters. Differentially expressed genes were assessed using DESeq2 version 1.28.1 (59). The Generalized Linear Model was fitted with the formula “design ≅ pairing + condition” to account for the pair-wise experimental design (control and treatment) whenever possible. Genes were considered differently expressed if they passed a threshold of adjusted P-value <0.05 (Benjamini-Hochberg correction) and |fold-change| >1.5.

### Identification and selection of reference genes

From the DESeq2 analysis of RNA-seq datasets, genes not differentially expressed between paired conditions were selected. TPM values were used to evaluate intra- and inter-experimental variability by calculating the coefficient of variation (CV) of each gene for each group and condition. The CV is defined by the ratio of the standard deviation of a gene TPM to the arithmetic mean of the same gene. Next, we calculated the arithmetic mean of CV between different conditions. Genes with a mean CV <0.10 (10%) were selected for further analysis. TPM values from the different conditions were averaged and the mean CV was evaluated within three subgroups of expression, namely genes with TPM between 10-100, 100-1000 and 1000-10000.

### Functional annotation

Functional enrichment was performed in R using clusterProfiler (60) and enrichplot (61) packages for Gene Ontology. The genes identified by RNA-seq in EndoC-βH1 cells and human islets with TPM >0.5 under control condition or after treatment were considered as expressed and used as background. An adjusted P-value <0.05 (Benjamini-Hochberg correction) was considered statistically significant.

### mRNA extraction, cDNA synthesis and quantification by qPCR

Poly(A)+ mRNA was isolated using Dynabeads mRNA Direct kit (Invitrogen), following the manufacturer’s protocol. mRNA was reverse transcribed using the Reverse Transcriptase Core kit (Eurogentec). In order to reduce variability in mRNA input, cDNA was quantified using the NanoDrop spectrophotometer (NanoDrop ND-1000; Thermo Fisher Scientific). Samples were then diluted to the same concentration of the least-concentrated sample (EndoC-βH1 cells were diluted to 800 ng/uL; dispersed human islets to 500 ng/uL; iPSC-beta cells to 300 ng/uL).

Primers were designed using Primer-BLAST software (62), using the following criteria: (I) primers were designed across exon-exon junction whenever possible; (II) PCR amplicon size of 80 −120 base pairs to minimize unwanted effects on the amplification efficiency; (III) primer Tm (melting temperature) was 56-60°C (with an annealing temperature of approximately 58°C); (IV) primer GC content was 40-65%, with no dimer or hairpin secondary structures. The qPCR amplification was done using IQ SYBR Green Supermix (Bio-Rad) using the CFX Connect Real-Time PCR Detection System (Bio-Rad). The amplification efficiency of each primer pair was evaluated using a standard curve (63), generated from six dilutions of cDNA (10^7^ to 10^2^ mRNA copies per microliter (copies per μL)). The target gene concentration was expressed as copies per μL (63). The Cq values specify the number of amplification cycles needed to detect a signal. Because Cq values are inversely correlated with the amount of target nucleotides, they were used to evaluate gene expression and extrapolate gene expression variability.

The cycling conditions were 95°C from 3 minutes (min), followed by 40 cycles of 95°C for 15 seconds (sec), and 58°C for 20 sec, followed by a final step of 95°C for 1min, 70°C for 5 sec, and 95°C for 50 sec. For each gene, the melting curve was analyzed to confirm amplification of a single PCR product. Primers are listed in **Supplementary table 1.**

### Statistical analysis

Significant differences between experimental conditions were determined by Student’s paired t-test or by ANOVA followed by Bonferroni correction as indicated. P-values <0.05 were considered statistically significant. Statistical tools used for RNA-seq analysis are described above. Violin plots illustrate kernel probability density (i.e., the width of the shaded area represents the data’s density plot). The horizontal line represents the median, boxes quartiles, whiskers most extreme data values, and dots independent experiments.

## Data accessibility

All raw RNA-sequencing data are accessible via NCBI Gene Expression Omnibus (GEO), access numbers GSE133218, GSE148058, GSE121863, GSE53949 and GSE159984.

---

**Supplementary Figure 1: Expression of new reference genes in beta cells from diabetic patients and validation by qPCR in EndoC-βH1 cells.** The newly identified reference genes *ARF1*, *CWC15*, *RAB7A*, *SIAH1*, *VAPA* are expressed in FACS-purified beta cells isolated from T1D patients **(A)** and islets from T2D patients **(B)**. Distribution of the quantification cycle (Cq) value of *ARF1*, *CWC15*, *RAB7A*, *SIAH1*, *VAPA*, *ACTB* and *GAPDH* based on qPCR in EndoC-βH1 cells treated with thapsigargin for 48h **(C)**, and *CWC15, RAB7A, VAPA* and *ACTB* in EndoC-βH1 treated with tunicamycin for 24h **(D).** For all samples the geometric mean of *ACTB* and *VAPA* Cq values was calculated (GM *ACTB VAPA*). The shape of the violin plots reflects the distribution of the data, and the width of curve corresponds to the frequency of the values in each region. The integrated boxplots show the 25^th^ and 75^th^ percentiles, and the horizontal line represents the median. Results from six to eight independent experiments. There is no statistical difference between the tested conditions (paired Student’s t-test, condition *versus* control (CTL) or non-treated (NT)).

**Supplementary Table 1:** Sequence of qPCR primers and siRNAs.

**Supplementary Table 2: Stably expressed genes in EndoC-βH1 cells and dispersed human islets under pro-inflammatory and metabolic stress conditions.** Coefficient of variance and transcripts per million (TPM) for 264 stably expressed genes with an adjusted P value >0.05, |fold change <1.5|.

## ACKNOWLEDGEMENTS

The authors are grateful to Isabelle Millard, Anyishaï Musuaya, Nathalie Pachera, Cai Ying, and Manon Depessemier (ULB Center for Diabetes Research) for providing excellent technical support.

