## Supplementary figures and images for "A functional genomic approach to identify reference genes for human pancreatic beta cell real-time quantitative RT-PCR analysis"

### Supplemental Figure 1

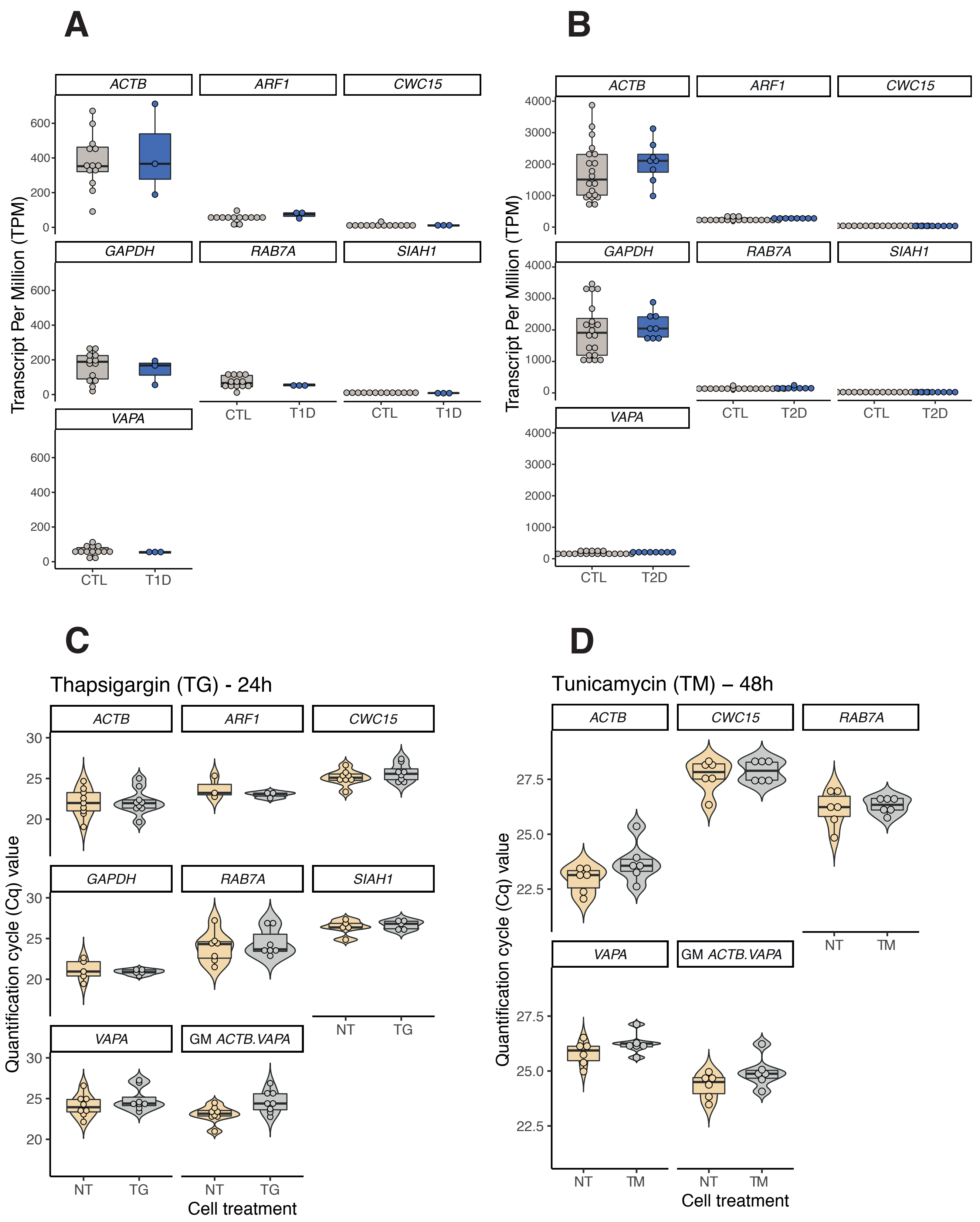
